# Thermoregulatory consequences of growing up during a heatwave or a cold snap

**DOI:** 10.1101/2023.04.21.537787

**Authors:** Elin Persson, Ciarán Ó Cuív, Andreas Nord

**Author notes:** Author for correspondence: Elin Persson, ORCID numbers: Elin Persson: 0000-0003-1824-1403 Ciarán Ó Cuív: 0009-0006-2958-8911 Andreas Nord: 0000-0001-6170-689X. AUTHOR CONTRIBUTIONS: Andreas Nord conceived the idea. Elin Persson, Ciarán Ó Cuív and Andreas Nord performed the practical work. Elin Persson analysed the data and all authors interpreted the results. Elin Persson wrote the first draft of the manuscript, which was edited by Andreas Nord and Ciarán Ó Cuív. All authors approved submission and agreed to be accountable for all contents. Andreas Nord procured funding. DATA AVAILABILITY: Data are deposited in Figshare: https://doi.org/10.6084/m9.figshare.24288067.v1. CONFLICT OF INTEREST: We have no conflicts of interest to disclose. SUMMARY STATEMENT: Early-life heatwaves or cold snaps affects thermoregulatory responses in Japanese quail chicks. However, no effects remained in adulthood once the extreme weather event had passed.

## Abstract

Changes in environmental temperature during the developmental period can affect growth, metabolism, and temperature tolerance of the offspring. We know little about whether such changes remain to adulthood, which is important to understand the links between climate change, development, and fitness. We investigated if phenotypic consequences of the thermal environment in early life remained in adulthood in two studies on Japanese quail (*Coturnix japonica*). Birds were raised under simulated heatwave-, cold snap-or control conditions, from hatching until halfway through the growth period, and then in a common garden until reproductively mature. We measured biometric and thermoregulatory (metabolic heat production [MHP], evaporative water and heat loss [EWL, EHL] and body temperature) responses to variation in submaximal air temperature at the end of the thermal acclimation period and in adulthood. Warm birds had lower MHP than control birds at the end of the thermal acclimation period and, in the warmest temperature studied (40°C), also had higher evaporative cooling capacity compared to controls. No analogous responses were recorded in cold birds, though they had higher EWL than controls in all but the highest test temperature. None of the effects found at the end of the heatwave-or cold snap period remained until adulthood. This implies that chicks exposed to higher temperatures could be more prepared to counter heat stress as juveniles, but that they do not enjoy any advantages of such developmental conditions when facing high temperatures as adults. Conversely, cold temperature does not seem to confer any priming effects in adolescence.

## Introduction

Climate change, including increased global mean temperature and increased climatic instability, has strongly impacted animal populations, for example, through changes in species distributions’ (Chen et al., 2011), mass mortality events (McKechnie et al., 2012; Moreno et al., 2015) and population extinctions (Thomas et al., 2006). Climate change is also associated with a range of sublethal consequences that can negatively affect fitness (Conradie et al., 2019). For example, many insectivorous passerines in the Northern Hemisphere have advanced breeding start to match a warming-related advancement of peak food abundance, but still pay fitness costs since phenotypic plasticity in egg laying cannot fully compensate for the earlier emergence of insect prey (Shipley et al., 2020; Both et al., 2009; Visser et al., 1998; Hansson et al., 2014). Climate change is further associated with increasingly variable environmental temperature. This can immediately impact growth and development, such as when cold snaps immediately reduce insect availability (Garrett et al., 2022; Sambaraju et al., 2012). Moreover, in response to acute and pronounced warming, such as during a heatwave, parents might have to trade-off workload against the risk of overheating (Cunningham et al., 2013; du Plessis et al., 2012; Nilsson and Nord, 2018) with implications for both current and future reproduction (Sharpe et al., 2019; Cunningham et al., 2021; Nord and Nilsson, 2019; Andreasson et al., 2020). Understanding the ontogeny and evolution of animals’ ability to cope with fluctuating environmental temperatures is thus fundamental to predict the consequences of climate change on wildlife (McKechnie and Wolf, 2019; Albright et al., 2017).

Changes in environmental temperature during the developmental period before or after hatching can affect subsequent temperature tolerance (reviewed by Nord and Giroud, 2020). Conceptually, it can be hypothesised that programming effects of developmental temperature on adult thermoregulation fall on a scale between constraining and priming: 1) if environmental temperature early in life programs phenotypes that perform optimally in that thermal environment, adults should maximise fitness when early-and later-life environments are similar (“environmental matching hypothesis”; reviewed by Monaghan, 2008); 2) if deviations from optimal developmental temperature act suppressively on the phenotype, then increasing distance from optimality in early life should come at a fitness cost regardless of the adult thermal environment (“silver spoon hypothesis”; Monaghan, 2008). In line with the environmental matching hypothesis, rats reared in warmer conditions grew their tails longer and feet larger and reduced the amount of body fat, which was interpreted as adaptions to facilitate heat loss (Demicka and Caputa, 1993). Also in line with this hypothesis, mice reared in colder conditions grew shorter tails and ears compared to mice reared in warmer conditions (Ballinger and Nachman, 2022) and cold exposed great tits (*Parus major*) grew smaller tarsi than control birds (Rodriguez and Barba, 2016a), which could be associated with reduced heat loss rate. Moreover, cold acclimated Pekin ducks (*Anas platyrhynchos*) and Japanese quail (*Coturnix japonica*) grew larger than control birds (Marjoniemi and Hohtola, 2000) with possible effects on both thermogenic capacity and thermal conductance. However, in wild systems, exposure to warmer developmental temperatures have been shown to have negative effects on nestling growth and fledging success. For example, several studies show that heat-stressed chicks grew more slowly than individuals reared in control conditions (Deaton et al., 1978; Andreasson et al., 2018; Rodriguez and Barba, 2016b; Cunningham et al., 2013; van de Ven et al., 2020; Andrew et al., 2017). These findings conform more closely to the silver spoon hypothesis. Other wild studies demonstrate positive effects of warmer developmental temperatures on nestling growth and fledgling success (McCarty and Winkler, 1999; Dawson et al., 2005). For example, tree swallow (*Tachycineta bicolor*) nestlings gained more mass due to higher environmental temperatures (McCarty and Winkler, 1999), and both nestling survival and fledging success increased with higher environmental temperatures in another study on the same species (Dawson et al., 2005). Moreover, studies show negative effects of exposure to colder developmental environmental temperatures on growth rate (Teulier et al., 2014), body mass (Ardia et al., 2010) and fledgling success (Garrett et al., 2022). This is not expected under the environmental matching hypothesis where lower body mass in the warmth and larger body mass in the cold should offer thermogenic advantage.

We know little about the physiological basis of any matching between developmental and adult thermal environments (reviewed by Nord and Giroud, 2020). Studies on poultry suggest that precisely timed and dosed thermal stimuli during embryonic or perinatal development can lead to epigenetic programming that makes the birds better at withstanding matched environmental temperature stressors, such as altered body temperature (*T*_b_) and metabolic rate, that last at least until adolescence (Nichelmann, 2004; reviewed by e.g. Farag and Alagawany, 2018). There are few comparable studies in wild systems. However, Andreasson et al. (2018) found that blue tit (*Cyanistes caeruleus*) nestlings from heated nest boxes had higher *T*_b_ and grew more slowly than nestlings from control nest boxes, indicating a constraining role of high environmental temperatures acting through a change in distribution of resources between growth and thermoregulation. Yet, heat-challenged nestlings in the study by Andreasson et al. (2018) tended to survive better to the next breeding season compared to control birds, providing mixed evidence for the constraining or priming role of developmental environmental temperature on adult phenotype. Furthermore, Page et al. (2022) found that blue tits that were experimentally heated as embryos were more cold-tolerant as mature nestlings than nestlings in non-heated nests. The above discrepancies suggest we need to know more about how morphological and physiological responses to developmental environmental temperature affect the capacity for thermal acclimation in a broader range of circumstances, particularly with regards to whether any of the reported phenotypic changes remain in adulthood. To fill this knowledge gap, we must measure the ontogeny and legacy of responses to variation in developmental environmental temperature, including changes in metabolic rate, evaporative cooling capacity and *T*_b_, both in the juvenile stage and in adulthood.

Here, we investigated if development during a simulated heatwave or cold snap affected offspring thermal, metabolic and hygric physiology, and if any such effects remained in adulthood several weeks after the heatwave or cold snap had passed. We studied this in Japanese quail that completed the first half of their somatic development under simulated heatwave-, or cold snap-like conditions or under normal thermal conditions, and the second half in a common garden. Epigenetic adaptation to environmental temperature is thought to appear most prominently during critical phases of development (Yahav, 2015). Even though precocial species like Japanese quail are covered with feathers and can thermoregulate from an early age, they are typically not fully homeothermic until some weeks after hatch (Jørgensen and Blix, 1988; Nord and Nilsson, 2021; reviewed by Price and Dzialowski, 2018). Accordingly, incipient thermogenesis develops from late incubation and over the first 10 or so days post-hatch (Yahav, 2015), suggesting there could be lasting effects on the thermoregulatory phenotype if chicks complete the first part of their post-embryonic life in heatwave or cold snap conditions. If warm developmental conditions prime heat acclimation, we predicted, under the environmental matching hypothesis, that heatwave birds would have lower *T*_b_ and correspondingly lower metabolic heat production rate (MHP) and a higher capacity for evaporative cooling during heat stress (Fig. 1A). If cold developmental conditions prime cold acclimation we predicted that cold snap birds would have higher *T*_b_ and higher MHP that render them better equipped to deal with low ambient temperature but, because of these adaptations, less efficient at evaporative cooling during heat stress (Fig. 1A). However, if high developmental ambient temperature constrains development, we predicted, under the silver spoon hypothesis, that heatwave birds would have higher *T*_b_, higher MHP and lower evaporative cooling capacity during a heat challenge (Fig. 1B). Cold snap birds would have lower MHP and lower *T*_b_ during a cold challenge (Fig. 1B). If ambient temperatures during the early developmental period permanently modifies temperature tolerance, as shown in studies on poultry (e.g., Arjona et al., 1988; Piestun et al., 2011), we also predicted that these phenotypic traits would remain in adulthood (Figs. 1A, B). If, however, developmental ambient temperature-effects are reversible, we predict that the change in traits caused by thermal acclimation would return to pre-acclimation levels according to the phenotypic flexibility hypothesis (Fig. 1C). We also measured body mass and wing length development, because in line with Bergmann’s rule, birds are predicted to grow larger when it is colder (Ashton, 2002) and smaller when it is warmer (Van Buskirk et al., 2010; Teplitsky et al., 2008). This is expected to decrease or increase dry heat loss rate due to a higher ratio between body surface area and volume. If a higher surface area to volume ratio is biologically meaningful to increase heat transfer rate in the warmth, we predicted that birds growing up under heatwave-like conditions would have lower body mass whereas birds growing up under cold snap-like conditions would have higher body mass in keeping with established ecogeographic rules (Ryding et al., 2021). This study provides new insights into how the developmental period influences how birds cope with increasing ambient temperatures in the short-and long term, by providing data on whether phenotypic effects are lasting, and under which circumstances such responses may be adaptive or maladaptive.

**Figure 1.**
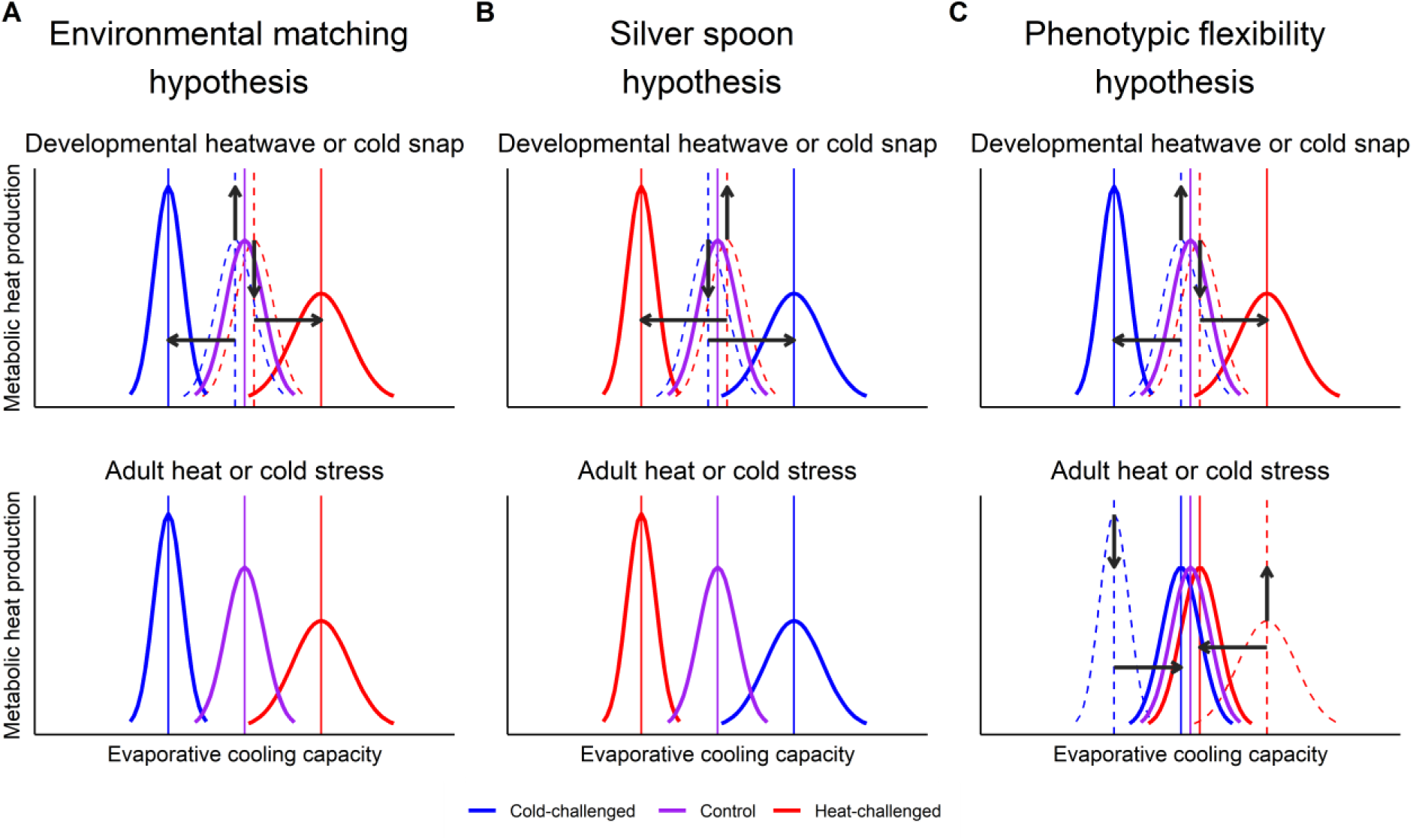
Three hypothetical scenarios for thermo-physiological changes in response to a developmental heatwave or cold snap. A) Under the Environmental matching hypothesis, a developmental heatwave leads to a non-reversible reduction in metabolic heat production and improvement in evaporative cooling capacity, such that individuals developing under heatwave-like conditions are more heat tolerant as adults whereas individuals developing under cold snap-like conditions are more cold tolerant as adults. B) Under the Silver spoon hypothesis, a change in environmental temperature during development impairs normal growth and maturation of the thermoregulatory system which lowers heat tolerance in heat-acclimated individuals and lowers cold tolerance in cold-acclimated individuals both in juveniles and adults. C) Under the Phenotypic flexibility hypothesis, a developmental heatwave triggers reduced metabolic heat production and improved evaporative cooling capacity and a developmental cold snap triggers increased metabolic heat production and reduced evaporative cooling capacity. However, these effects are reversible so that the developmental thermal environment does not affect temperature tolerance in adulthood. Arrows show predicted direction of changes. Dashed lines indicate trait starting points.

## Methods

### Species and husbandry

The study was performed in two iterations: acclimation to heatwave-like conditions (henceforth: Warm study) was performed between April and July 2021 whereas acclimation to cold snap-like conditions (henceforth Cold study) was performed between September and December 2021. Both studies followed the same general design.

Japanese quail eggs (Warm study: n=60; Cold study n=92) were bought from a commercial breeder (Sigvard Månsgård, Åstorp, Sweden) and were incubated in 37.9 ± 0.52°C at 50% relative humidity using Brinsea OvaEasy 190 incubators (Brinsea, Weston-super-Mare, United Kingdom). The eggs were stored at room temperature (18-20°C) and were turned manually twice daily (Warm study) or automatically for 5 min every 1 h (Cold study) until put into the incubator (6 eggs per day). On day 15 of incubation, the eggs were moved from the incubation trays to a hatching tray contained inside the incubator. The eggs were checked for signs of hatching daily from day 17 onwards. The incubation period was 19 days (range: 17 to 21d) in the warm study and 18 days (range: 17 to 20d) in the cold study. In the Warm study, 45 of 60 eggs (75%) hatched, in the Cold study, 47 of 92 eggs (51%) hatched. The lower hatching success in the Cold study coincided with a drop in fertility reported by the breeder.

After hatching, the chicks were left inside the incubator until completely dry or for a maximum of 12 hours. They were then transferred to open pens (310 × 120 × 60 cm) dressed with wood shavings. During the first 3 weeks after hatching, housing temperature was either 20°C (henceforth “control treatment”; Warm and Cold study), 30°C to simulate heatwave conditions (henceforth “warm treatment”) in the Warm study, or 10°C to simulate cold snap conditions (henceforth “cold treatment”) in the Cold study. From then on, all chicks were transferred to new pens and were housed in a common garden at 20°C until the end of the experiment. These temperatures were near the upper and lower critical temperatures, respectively, in Japanese quail chicks (Ben-Hamo et al. 2010) and broadly similar to those used by others studying the thermal sensitivity of development in quail (Burness et al., 2013). Chicks were assigned randomly to the warm, cold or control treatments. Final samples sizes are reported in Table 1. The quail had *ad libitum* access to pelleted feed, water, sandbathes and crushed seashells throughout the experiment. Mealworms and vegetables (kale, carrot, or lettuce) were each offered once weekly on alternate days. Until 3 weeks old, the birds were fed turkey starter (Kalkonfoder Start, Lantmännen, Stockholm, Sweden; 25.5% protein). After 3 weeks of age, the feed was switched to turkey grower (Kalkonfoder Tillväxt, Lantmännen, Stockholm, Sweden; 22.5% protein). Until 3 weeks old, all birds had access to a heat lamp providing an ambient temperature of 37°C at floor level, but no heating source was provided from week 3 onwards. The heat lamp was placed so that the quail had to experience experimental room temperatures to access feed, water and enrichments. Photoperiod was 12:12 LD (Warm study: lights on from 06:00 to 18:00 GMT+2; Cold study: lights on from 07:00 to 19:00 GMT+1) throughout the experiment.

**Table 1.** The number of Japanese quail used when studying morphological and thermo-physiological responses of developing in either 30°C (Warm study), 20°C (control; both studies) or 10°C (Cold study) for the first 3 weeks of life, and in 20°C thereafter. Morphological responses (body mass, wing length) were measured once weekly, except at 4 and 6 weeks of age. Thermo-physiological responses (metabolic heat production (MHP), evaporative water loss (EWL), body temperature (*T*_b_)) were measured at increasing air temperature at the end of the temperature treatment, and after 5 weeks in a common garden. The same birds were measured at all ages, and at all temperatures, but some experiments and measurements were excluded from the final data set for reasons stated in the main text. The starting number of quail in the morphological analyses is stated under *n*_biometry_, whereas the total number of birds included in the thermal and metabolic analyses are stated under *n*_respirometry_.

| | $n_{\text{biometry}}$ | $n_{\text{respirometry}}$ | Sample size | | | |
| --- | --- | --- | --- | --- | --- | --- |
|  |  |  | 10°C | 20°C | 30°C | 40°C |
| **Warm study** | **44** |  |  |  |  |  |
| <u>3 weeks</u> |  | 39 |  |  |  |  |
| Control (6 ♀, 10 ♂, 2 unknown) |  |  | 17 | 17 | 17 | 15 |
| Warm (12 ♀, 5 ♂, 5 unknown) |  |  | 22 | 22 | 22 | 19 |
| <u>8 weeks</u> |  | 36 |  |  |  |  |
| Control (6 ♀, 10 ♂, 1 unknown) |  |  | 16 | 15 | 16 | 15 |
| Warm (12 ♀, 5 ♂, 3 unknown) |  |  | 18 | 19 | 19 | 18 |
| **Cold study** | **34** |  |  |  |  |  |
| <u>3 weeks</u> |  | 36 |  |  |  |  |
| Control (11 ♀, 7 ♂) |  |  | 16 | 17 | 18 | 16 |
| Cold (10 ♀, 8 ♂) |  |  | 18 | 18 | 17 | 18 |
| <u>8 weeks</u> |  | 35 |  |  |  |  |
| Control (11 ♀, 6 ♂) |  |  | 16 | 16 | 16 | 16 |
| Cold (11 ♀, 7 ♂) |  |  | 15 | 17 | 15 | 17 |

### Measurements of body size and body temperature

Morphometric measurements were taken starting on day 1 and then once weekly throughout the experiment. On day 1, birds were banded and weighed (± 0.1 g). From day 7 onwards, wing length was also measured (± 0.5 mm). When the birds were 13 to 16 days old, we implanted a temperature-sensitive PIT tag (LifeChip BioTherm, Destron Fearing, South St. Paul, MN, USA), 2.1 × 12 mm in size (< 0.5% of body weight), into the intraperitoneal cavity to measure *T*_b_. Ventral skin surrounding the distal end of the sternum was sterilised (70% EtOH) and anaesthetized topically using 5% lidocaine (EMLA*®*, Aspen Pharma, Durban, South Africa). Thirty to 45 min later, the skin was sterilized again, and a sterile PIT tag was inserted immediately beneath the distal tip of the sternal keel using a sterile 12-gauge syringe. This positioned the PIT tag on top of the liver (Andreas Nord, pers. obs.). The incision was closed using UHU cyanoacrylate (Bolton Adhesives, Rotterdam, Nederland) and covered with antiseptic ointment (1% H_2_O_2_, LHP, Bioglan AB, Malmö, Sweden). We calibrated the PIT tag antennae at 35, 40, and 45°C using a subset of 12 tags (i.e., 3 tags for each of the 4 antennae used in the Studies) and used the antenna-specific calibration equations to correct *T*_b_ data before further analyses.

### Measurements of metabolic heat production and evaporative water loss

Metabolic heat production (MHP) and evaporative water loss (EWL) were measured using flow-through respirometry in a climate test chamber (Weiss Umwelttechnik C180, Reiskirchen, Germany). Measurements were taken, firstly, at the end of the heatwave-or cold snap treatment when the birds were 3 weeks old and halfway through the somatic growth phase. Measurements were taken again at 8 weeks, when the birds had reached asymptotic size and started reproducing (i.e., when they were young adults). During the first experimental period, we used 3.3 L glass respirometry chambers, and for the second experimental period 8.0 L glass chambers. The chambers were ventilated with dry (drierite; Sigma-Aldrich, Stockholm, Sweden) atmospheric air, in the Warm study at 1.873 – 2.638 L/min (mean ± s.e.m.: 2.149 ± 0.013 L/min; standard temperature and pressure, dry, STPD) and in the Cold study at 2.047 – 2.272 L/min (mean ± s.e.m.: 2.152 ± 0.003 L/min; STPD) during the 3 week measurements and 3.653 – 4.751 L/min (mean ± s.e.m.: 4.179 ± 0.013 L/min, STPD) in the Warm study and 4.093 – 4.558 L/min (mean ± s.e.m.: 4.313 ± 0.008 L/min, STPD) in the Cold study during the 8 week measurements as measured with a FB8 mass flow meter (Sable Systems, Las Vegas, NV, USA). The 99 % equilibrium times (Lasiewski et al., 1996) were 5.8 – 8.1 min and 7.8 – 10.1 min (Warm study at 3 and 8 weeks respectively), and 6.7 – 7.4 min and 8.1 – 9.0 min (Cold study at 3 and 8 weeks). We sub-sampled 308 – 388 ml/min (mean ± s.e.m.: 333 ± 1 ml/min, STPD) in the Warm study and 308 – 388 ml/min (mean ± s.e.m.: 369 ± 1 ml/min, STPD) in the Cold study from the main gas stream using a SS-4 sub-sampler (Sable Systems) for gas analyses. Oxygen was measured using a FC-10 analyser (Sable Systems) and water vapor was measured with a RH-300 water vapour meter (Sable Systems). Water vapour and carbon dioxide were removed from the airstream before measuring oxygen using drierite and ascarite (II) (Acros Organics, Geel, Belgium). The oxygen analyser was zero calibrated using 100% N_2_ and span calibrated to 20.95% O_2_ using dry (drierite) room air. The RH-300 was zero calibrated using dry room air, and span calibrated by exploiting the dilution of oxygen by water vapour in moist air. Thus, by recording barometric pressure (BNP, kPa) and O_2_ content in dry (O_2dry_) and wet (O_2wet_) air, it is possible to calculate water vapour pressure (WVP, kPa):

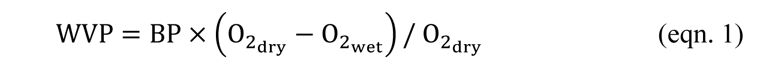

Birds were placed on a metal grid platform over a reservoir of mineral oil to avoid that evaporation from faeces affected the measurements of EWL (Lasiewski et al., 1966). Air temperature in the chambers were measured using thermocouples (36-gauge type T, copper-constantan) connected to a thermocouple box (TC-2000, Sable Systems) at a height where the temperature reading was not affected by contact with the bird. *T*_b_ was measured at 1 Hz throughout the experiment by placing custom-made antennas connected to a RM310 multiplexer board (Biomark, Boise, ID, USA) below the respirometry chambers. Measurements were performed with the climate test chamber set at 10, 20, 30 and 40°C. The corresponding air temperatures in the respirometer chambers were 12 ± 0.14, 21 ± 0.07, 31 ± 0.05 and 40 ± 0.07°C in the Warm study and 10 ± 0.18, 20 ± 0.11, 30 ± 0.05 and 40 ± 0.03°C in the Cold study.

All experiments took place during daytime. Birds were placed in a climate chamber illuminated by a light bulb at least 30 min before the start of data collection for thermal acclimation to the first measurement temperature (which was always 10°C). *T*_b_ initially rose when birds were handled for placement in into the respirometry chambers, but this handling-induced increase had worn off by the time the experiment had started (mean stabilising time ± s.e.m.: 13 ± 1 min). During the first experimental period, a measurement cycle started with a 10-min baseline followed by 10 min of sequential data collection for each of four birds and ended with a 20-min baseline (i.e., exposure: 70 min per measurement temperature and 280 min for a full experiment). During the second experimental period (i.e., when birds were 8 weeks old), 2 or 3 birds were measured simultaneously. Measurements started with a 15-min baseline followed by 10 min of data collection for each bird and ended with a 20-min baseline (i.e., exposure: 55 to 65 min per measurement temperature; 220 to 260 min per experiment).

Air temperature in the climate chamber was increased acutely to the next target measurement temperature at the start of the last baseline measurement in a cycle, the new target being reached within 15 min resulting in a minimum of 10 min of acclimation to the new target before data collection started again. A bird was removed from the experiment if it showed signs of distress that did not pass in 5 min, or if *T*_b_ rose > 45°C at the same time as it showed signs of distress.

### Data analyses

#### Missing data

Of the 45 birds that hatched in the Warm study, 39 (control: 17; warm: 22) were subsequently used in the first experimental period and 36 (control: 16; warm: 20) in the second experimental period (Table 1). Four birds died of natural causes or were euthanized before any measurements took place, and an additional 3 died between 3 and 8 weeks of age (before 5 weeks: 2; before 7 weeks: 1). Furthermore, 1 bird was never considered for any measurements because it showed abnormally slow growth rate and was subsequently subject to severe pecking by the other birds. One bird was left out of the thermal and metabolic experiments because it could not be fitted into the measurement schedule (see above) but was included in the analyses of morphological characteristics. Five birds were removed from the experiment before the measurements at 40°C at 3 weeks of age due to signs of stress (see above). Data for 8 birds from 6 sessions (of 100 total) were excluded from the data set because gas concentrations were not at steady state, typically because the birds were non-resting and so did not conform to the criteria for measuring resting metabolic rate (IUPS Thermal Commission, 2003). Data from 2 birds were excluded from the wing length (1 control, 5-week-old bird) and body mass (1 warm, 5-week-old bird) analyses due to measurement errors. Of the 47 birds that hatched in the Cold study, 11 died from natural causes or were euthanized before the first experimental period (not included in any analyses) and 1 bird died between 7 and 8 weeks of age. A total of 36 (control: 18; cold: 18) birds were measured in the first experimental period and 35 (control: 17; cold: 18) in the second experimental period (Table 1). Two birds were removed from the experiments before the measurement at 40°C at 3 weeks due to stress. Data from 7 birds from 10 sessions (of 116) were subsequently excluded from the data because the gas consumption curves were not at steady state. Two control birds were excluded from the morphological analyses. Wing length was not measured in 10 birds at 1 week of age (control: 5; cold: 5). A breakdown of sample sizes per ages and response variables in each Study is provided in Table 1.

#### Calculations

Data were extracted using ExpeData (version 1.9.27; Sable Systems). Oxygen consumption (ml / min) was calculated from the most stable 2 min period of the 10 min recording using eqn. 11.1 in Lighton (2008), and was converted to metabolic heat production (MHP, in W) assuming 1 ml of O_2_ = 20 J (Kleiber, 1961). Evaporative water loss (EWL, in mg / min) was calculated from the same period using eqn. 11.9 in Lighton (2008) and was converted to EHL (in W), assuming 2406 J is required to evaporate 1 ml of H_2_O (Wallace and Hobbs, 2006). We then calculated evaporative cooling efficiency as the ratio between EHL and MHP (Lasiewski et al., 1966). Body temperature was calculated as the mean of the 2 min period for which MHP and EWL were defined.

#### Statistical analyses

Data from the two studies were analysed separately. Statistical analyses were made in R (version 4.0.3; R Core Team, 2023) using linear mixed models (lmer function in lme4; Bates et al., 2015). To test if the treatment affected size and growth, we used body mass or wing length as dependent variables, treatment as a factor, and age and age^2^ as co-variates. The original model also included the interactions treatment × age and treatment × age^2^. Bird ID was used as a random factor to account for repeated measurements on the same individual. For the thermal and metabolic models, MHP, EWL, EHL/MHP, and *T*_b_ were used as response variables. To test the effects of measurement temperature on these metrics, we fitted separate models at 3 and 8 weeks of age with treatment, air temperature, and treatment × air temperature as factors, body mass as a continuous covariate and bird ID as random intercept (to account for repeated measurements). MHP, EWL and EHL/MHP were log-transformed before the analyses to meet parametric assumptions.

*P*-values for both fixed and random factors were assessed using likelihood ratio tests. The interaction was removed from the model when non-significant (i.e., *P* > 0.05), but all main effects were retained. If interactions were significant, a post hoc test (using the pairs() function in the emmeans package; Lenth, 2016) was performed between treatments within air temperatures. Model estimates where calculated using the emmean() function in the emmeans package (Lenth, 2016), and were back-transformed from the log scale when applicable. Figures show raw data.

## Results

### Body mass and wing length

In both studies, body mass and wing length increased non-linearly with age, with asymptotic phases for both traits reached between 5 and 6 weeks of age. Neither growth curve differed between the treatments. Nor was there any difference in the body mass (Fig. 2A; 2B) or wing length (Fig. 2C; 2D) intercepts between warm-and control birds in the Warm study, or between cold-and control birds in the Cold study (Table 2). Birds had reached 58% (Warm study) and 55% (Cold study) of their adult (i.e., 8 week) body mass, and 81% (Warm study) and 77% (Cold study) of adult wing length at 3 weeks of age (Table 2; Fig. 2).

**Figure 2.**
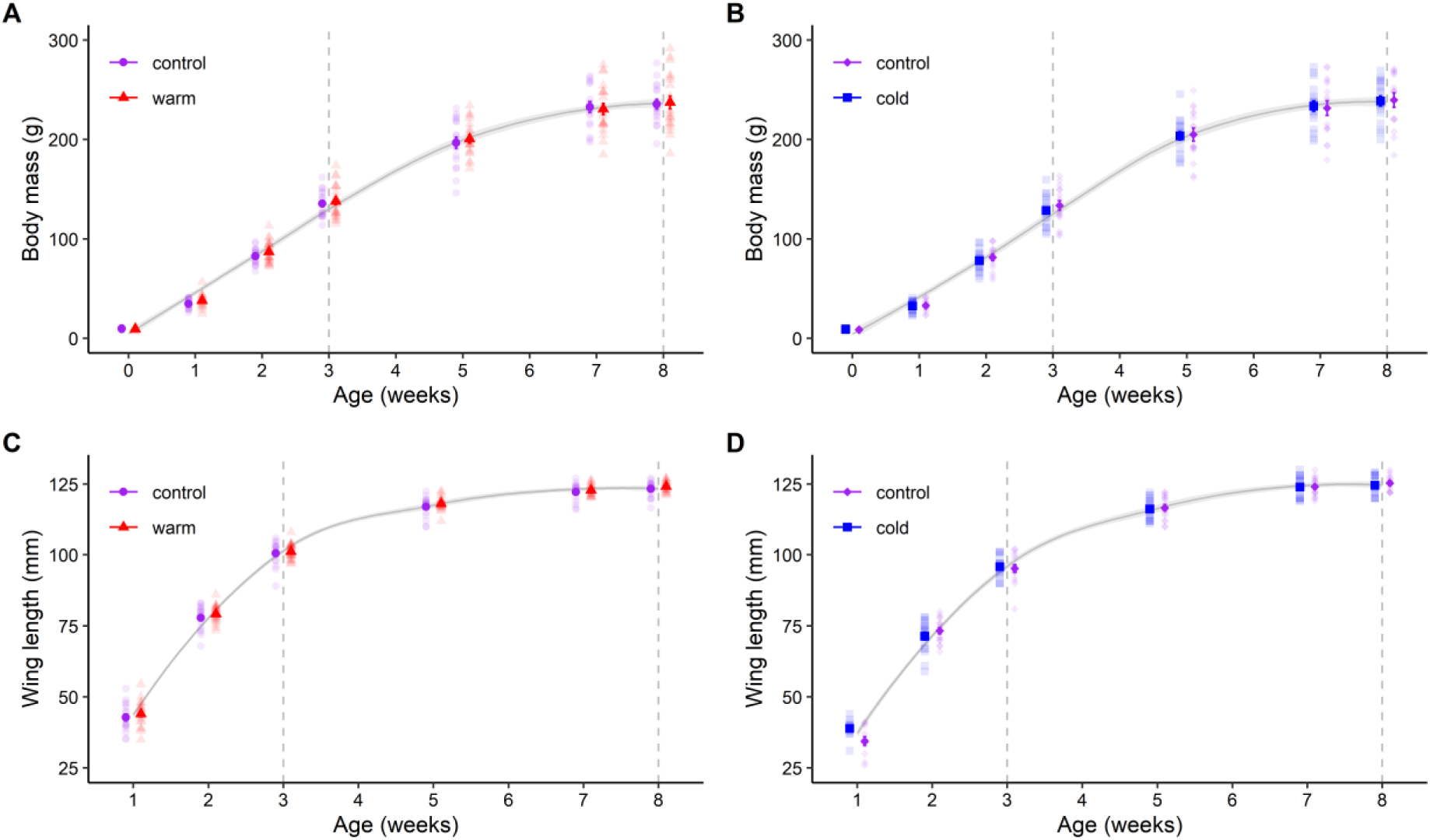
Effects on body mass and wing length when Japanese quail developed in either control (20°C), warm (30°C) or cold (10°C) temperature conditions. The figure shows mean ± s.e.m. (A: Warm study; B: Cold study) body mass and (C: Warm study; D: Cold study) wing length of Japanese quail during the 8-week experimental periods. The grey curve is LOESS (locally estimated scatterplot smoothing) ± 95% confidence interval. The dashed lines represent the timing of measurements of metabolic rate and evaporative water loss. Sample sizes per age are stated in Table 1. Semi-transparent points show raw data. Data were collected in two separate studies as detailed in the main text.

**Table 2.** Parameter estimates from linear mixed models explaining the effects of developing in either control (20°C) warm (30°C) or cold (10°C) temperature conditions in Japanese quail. Estimates, test statistics, degrees of freedom, P-values and standard deviations (σ) for the random factor on body mass and wing length. Data on warm-and cold-acclimation were collected in two sequential studies, which were analysed separately. Sample sizes are presented in Table 1.

| Model | Estimate ± s.e.m. | LR | Df | P | $\sigma_{\text{ID}}$ \| $\sigma_{\text{total}}$ |
| --- | --- | --- | --- | --- | --- |
| **Warm study** |  |  |  |  |  |
| <u>Body mass (g)</u> |  |  |  |  |  |
| Treatment |  | 0.19 | 1 | 0.6601 |  |
| Age | 58.40 ± 1.497 | 476.71 | 1 | <0.0001 *** | |
| Age<sup>2</sup> | -2.83 ± 0.145 | 225.62 | 1 | <0.0001 *** | |
| Treatment:Age |  | 0.30 | 1 | 0.5870 |  |
| Treatment:Age<sup>2</sup> |  | 0.39 | 1 | 0.5352 |  |
| Random factor bird ID |  | 67.01 | 1 | <0.0001 *** | 11.44 \| 25.69 |
| <u>Wing length (mm)</u> |  |  |  |  |  |
| Treatment |  | 2.07 | 1 | 0.1500 |  |
| Age | 35.07 ± 0.720 | 560.34 | 1 | <0.0001 *** | |
| Age<sup>2</sup> | -2.75 ± 0.078 | 429.19 | 1 | <0.0001 *** |  |
| Treatment:Age |  | 0.01 | 1 | 0.9417 |  |
| Treatment:Age<sup>2</sup> |  | 0.00 | 1 | 0.9776 |  |
| Random factor bird ID |  | 0.08 | 1 | 0.7844 | 0.60\|5.99 |
| **Cold study** |  |  |  |  |  |
| <u>Body mass (g)</u> |  |  |  |  |  |
| Treatment |  | 0.06 | 1 | 0.7998 |  |
| Age | 52.88 ± 1.423 | 419.78 | 1 | <0.0001 *** |  |
| Age<sup>2</sup> | -2.73 ± 0.170 | 167.84 | 1 | <0.0001 *** |  |
| Treatment:Age |  | 0.39 | 1 | 0.5315 |  |
| Treatment:Age<sup>2</sup> |  | 0.54 | 1 | 0.4614 |  |
| Random factor bird ID |  | 62.13 | 1 | <0.0001 *** | 12.88\|28.28 |
| <u>Wing length (mm)</u> |  |  |  |  |  |
| Treatment |  | 0.01 | 1 | 0.9332 |  |
| Age | 35.60 ± 1.541 | 474.65 | 1 | <0.0001 *** |  |
| Age<sup>2</sup> | -2.68 ± 0.077 | 358.70 | 1 | <0.0001 *** |  |
| Treatment:Age |  | 1.04 | 1 | 0.3084 |  |
| Treatment:Age<sup>2</sup> |  | 0.84 | 1 | 0.3599 |  |
| Random factor bird ID |  | 17.52 | 1 | <0.0001 *** | 2.70\|7.43 |

### Effects of air temperature on thermal and metabolic responses

There was no interaction between treatment and air temperature on log(MHP) at any age in neither the Warm nor the Cold study (Table 3). At 3 weeks, the warm-acclimated birds (i.e., in the Warm study) had a lower log(MHP) (1.88 ± 0.049 W) than control birds (2.03 ± 0.060 W) (Table 3; Fig. 3A), but there was no effect of treatment on log(MHP) at 8 weeks (Table 3; Fig. 3A). Log(MHP) was not affected by treatment at any age in the Cold study (Table 3; Fig. 3B). Log(MHP) increased with increasing body mass both at 3 and 8 weeks in the Warm study and at 3 weeks in the Cold study (Table 3). Also, in both studies, exposure to submaximal heat (i.e., 40°C) was associated with increased MHP at 3 weeks, but at 8 weeks MHP in the highest test temperature did not increase above minimal values.

**Figure 3.**
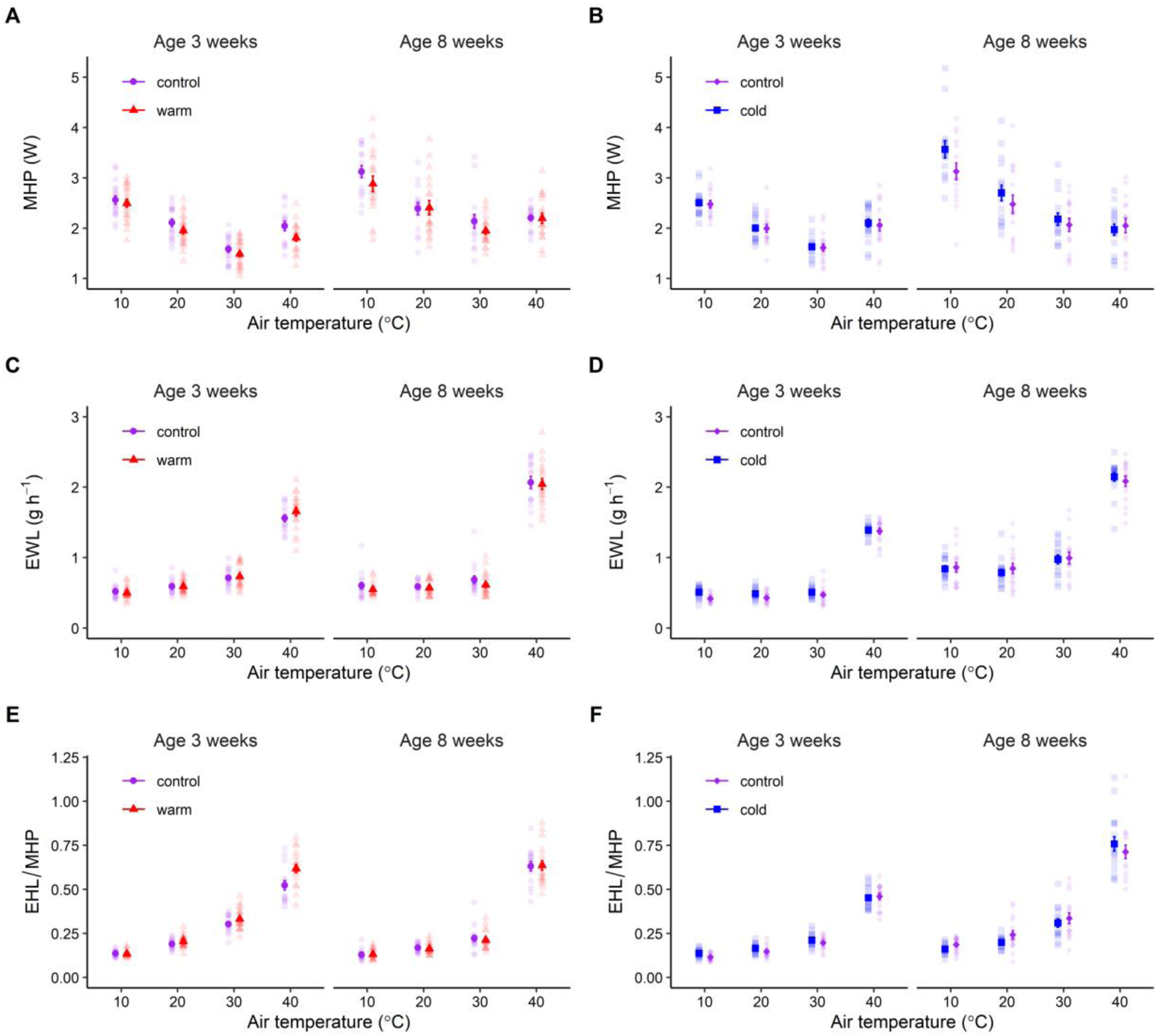
Effects on metabolic heat production (MHP), evaporative water loss (EWL) and evaporative cooling capacity (EHL/MHP) when Japanese quail developed in either control (20°C), warm (30°C) or cold (10°C) temperature conditions. The figure shows mean ± s.e.m. (A: Warm study; B: Cold study) MHP, (C: Warm study; D: Cold study) EWL and (E: Warm study; F: Cold study) EHL/MHP of Japanese quail in the different test temperatures (10°C, 20°C, 30°C and 40°C) at 3 weeks and 8 weeks of age. Sample sizes per age and temperature are stated in Table 1. Semi-transparent points show raw data. Warm-and cold-acclimation data were collected in two separate studies, as detailed in the main text.

**Table 3.**
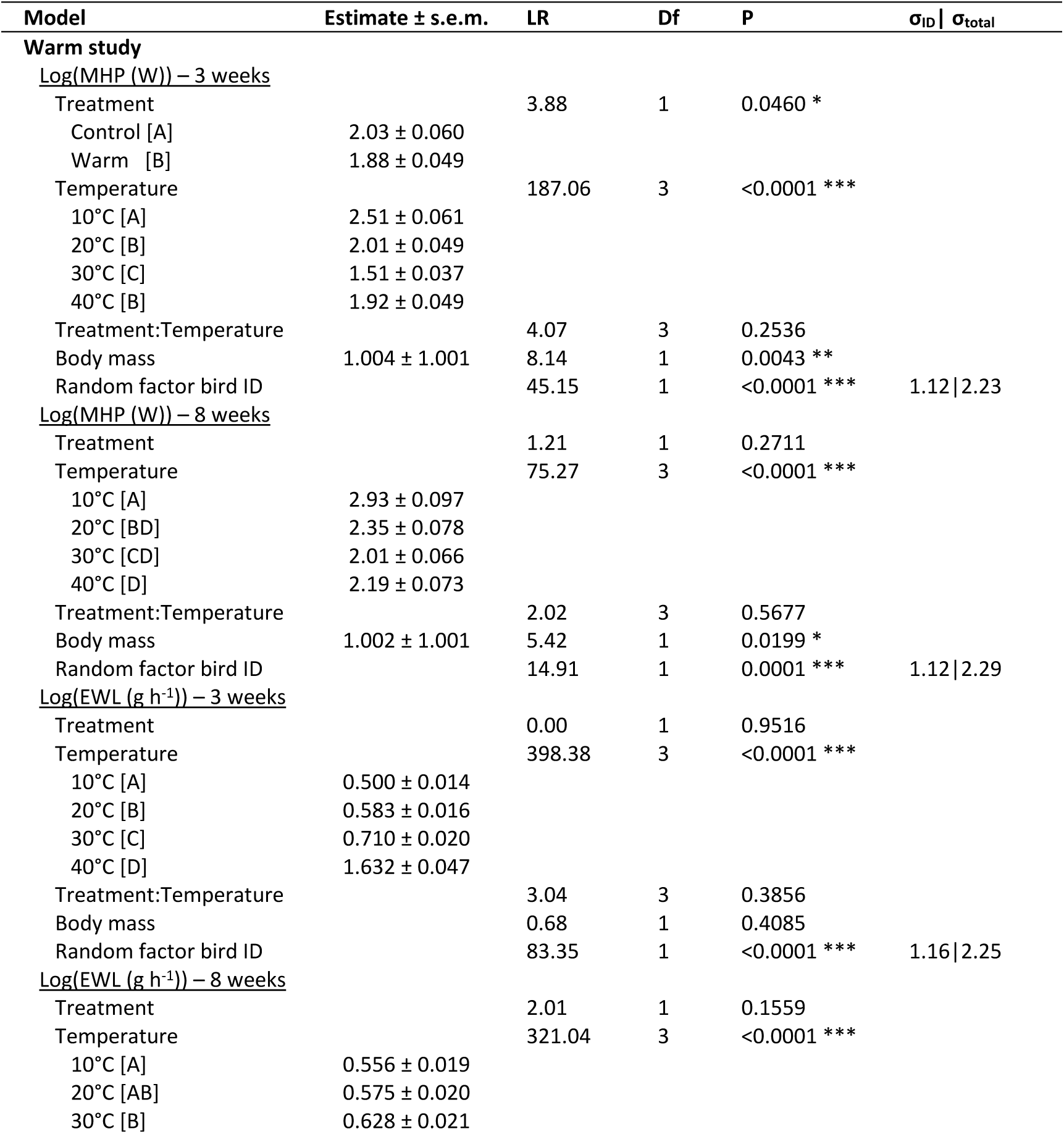
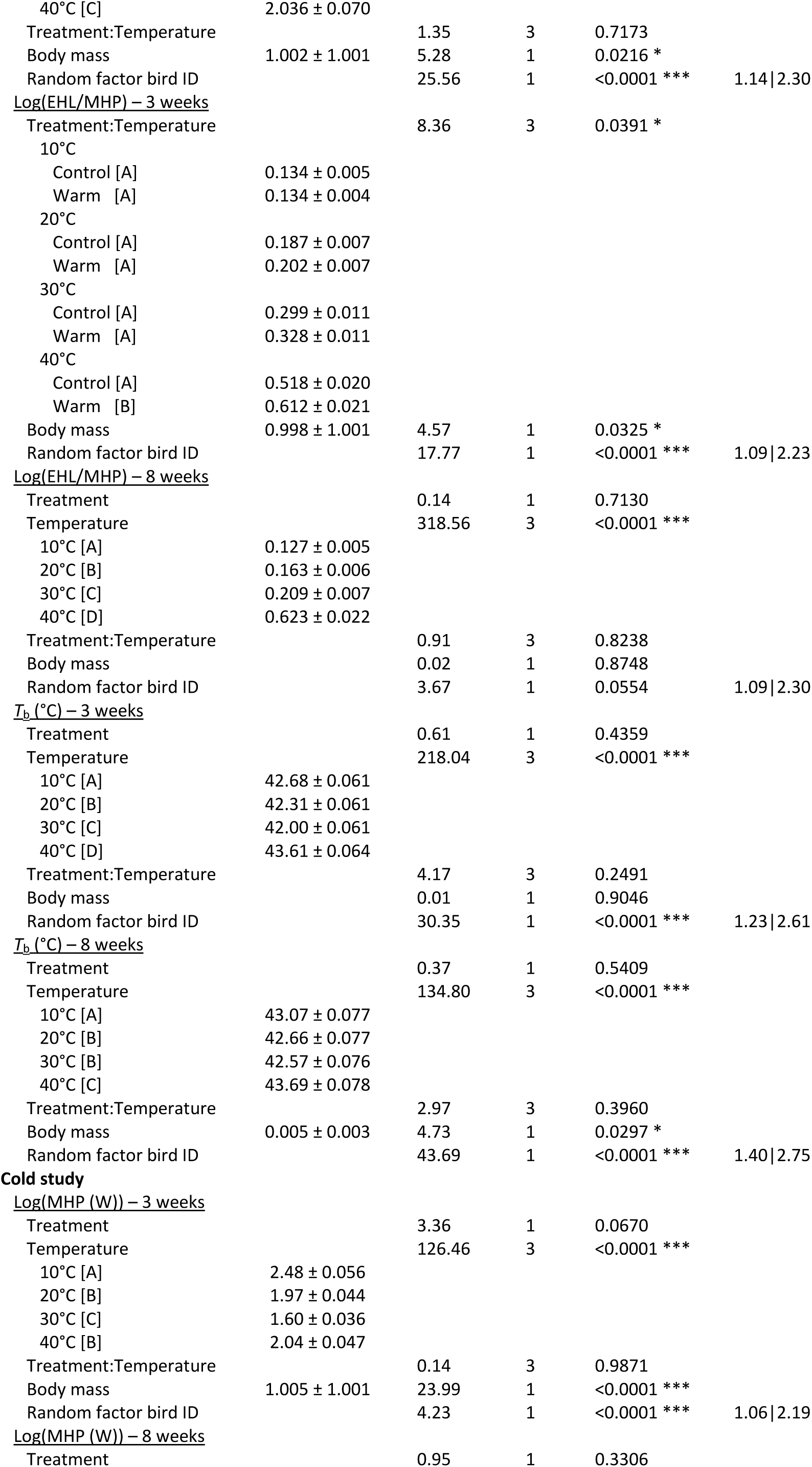
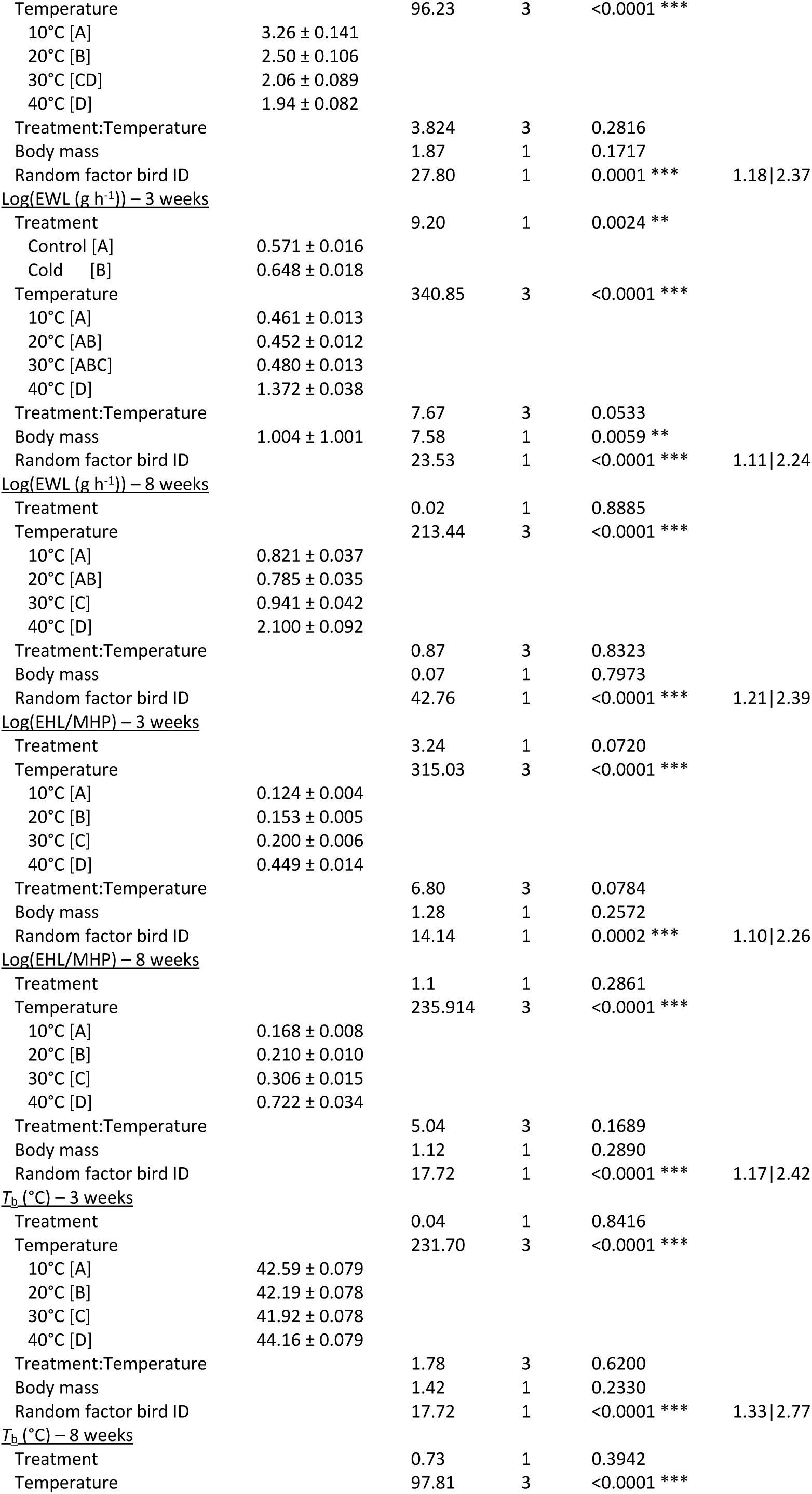
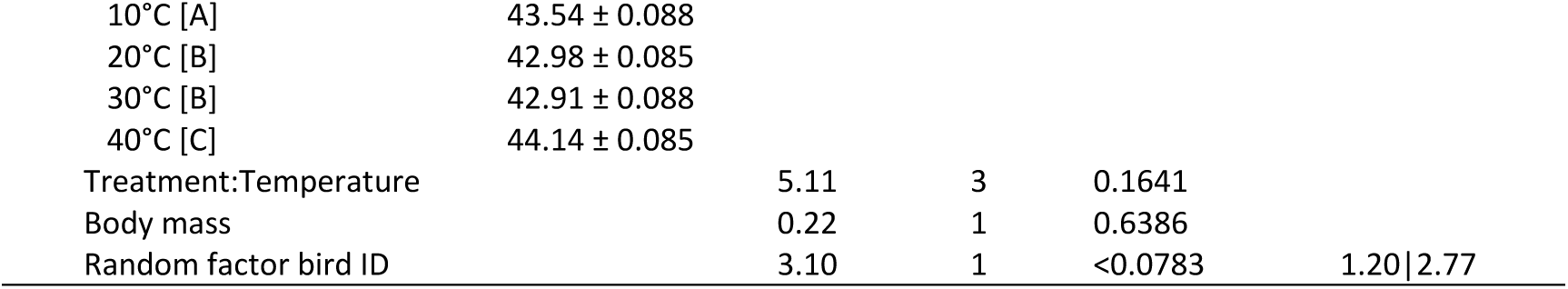
Parameter estimates from linear mixed models explaining the effects of developing in either warm (30°C), cold (10°C) or control (20°C) temperature conditions in Japanese quail. Estimates, test statistics, degrees of freedom, P-values and standard deviations (σ) for the random factor on treatment at 3-and 8 weeks on metabolic heat production, evaporative water loss, evaporative cooling capacity (i.e., evaporative heat loss/metabolic heat production) and body temperature. All dependent variables were log-transformed in the models, but back-transformed model estimates are shown in the table. Statistics for main effects is not provided when the interaction between treatment and temperature was significant. Different letters within brackets represent significant (P < 0.05) post hoc comparisons. Data on warm-and cold-acclimation were collected in two sequential studies, which were analysed separately. Sample sizes are presented in Table 1. Abbreviations: MHP: Metabolic heat production; EWL: Evaporative water loss; EHL/MHP: Evaporative heat loss/Metabolic heat production; *T*_b_: Body temperature.

Log(EWL) was not affected by the interaction between treatment and air temperature, nor by treatment alone at any age in the Warm study (Table 3). In the Cold study, the effect of treatment × air temperature was nearly significant (*P* = 0.053; Table 3). This interaction manifested as cold birds having higher log(EWL) in 10°C, 20°C and 30°C, but not at 40°C, compared to control birds. When the interaction was removed from the Cold study model, cold birds had higher log(EWL) (0.648 ± 0.018 g h^-1^) than control birds (0.571 ± 0.016 g h^-1^) (Table 3; Fig. 3D) across all air temperatures. At 3 and 8 weeks in both studies, log(EWL) increased with air temperature (Table 3; Fig. 3C; 3D). In the Warm study, log(EWL) was not affected by body mass at 3 weeks (Table 3), but it increased with body mass at 8 weeks (Table 3). In the Cold study, log(EWL) increased with body mass at 3 weeks, but not at 8 weeks (Table 3).

Evaporative cooling capacity which measures how large a proportion of MHP that the birds lost by evaporation (i.e., log(EHL/MHP)), was affected by the interaction between treatment and air temperature at 3 weeks in the Warm study. Specifically warm birds lost a larger proportion of MHP (0.612 ± 0.021) than control birds (0.518 ± 0.020) in 40°C (Table 3; Fig. 3E), but there was no difference between the treatments in any other air temperature (Table 3). At 3 weeks in the Cold study, log(EHL/MHP) was not affected by the interaction between treatment and air temperature or by the main effect of treatment, but it increased with air temperature (Table 3; Fig. 3F). At 8 weeks, log(EHL/MHP) was not influenced by treatment, neither as a main effect or when interacting with air temperature (Table 3) in either Study, but it increased significantly with air temperature (Table 3; Fig. 3E; 3F). At 3 weeks in the Warm study, log(EHL/MHP) decreased with increasing body mass (Table 3)but this effect had disappeared by week 8 (Table 3). In the Cold study, log(EHL/MHP) was not affected by body mass at any age (Table 3).

*T*_b_ did not differ between treatments at any age in either Study, neither when considering the main effect nor its interaction with air temperature (Table 3). However, *T*_b_ increased with both decreasing and increasing air temperature across treatments for both ages in both studies (Table 3; Fig. 4A; 4B). Higher *T*_b_ at lower air temperatures probably reflects the combined effects of thermogenesis and insulation whereas, by analogy, higher *T*_b_ in the warmth reflects insufficient heat loss. There was no effect of body mass on *T*_b_ at 3 weeks in either Study (Table 3). At 8 weeks, *T*_b_ increased with increasing body mass in the Warm study but not in the Cold study (Table 3).

**Figure 4.**
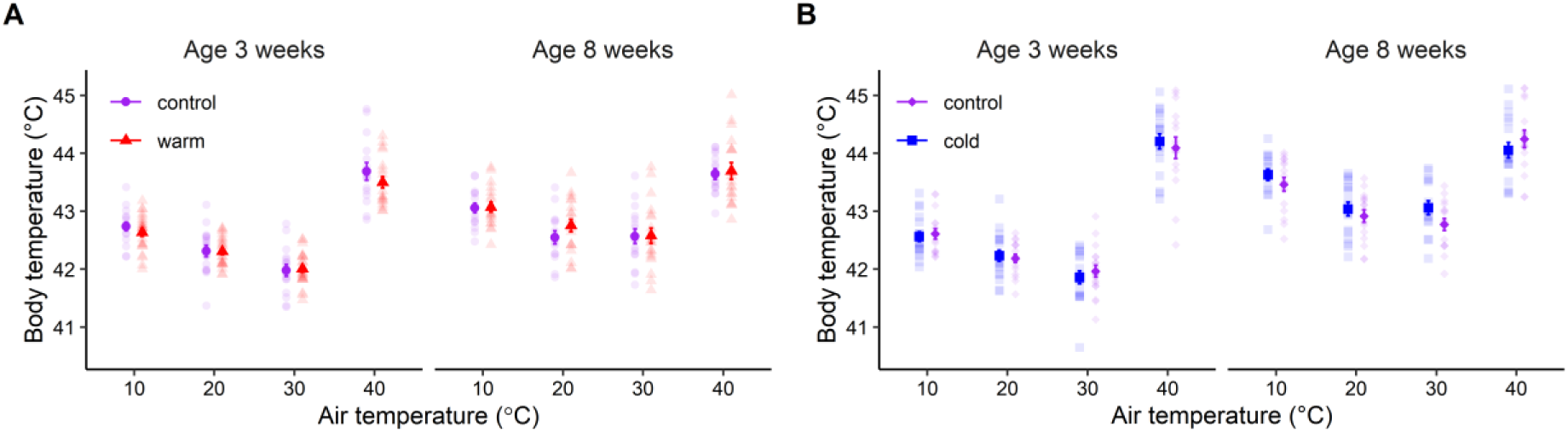
Effects on body temperature (*T*_b_) when Japanese quail developed in either control (20°C), warm (30°C) or cold (10°C) temperature conditions. The figure shows (A: Warm study; B: Cold study) mean ± s.e.m. *T*_b_ of Japanese quail in the different test temperatures (10°C, 20°C, 30°C and 40°C) at 3 weeks and 8 weeks of age. Sample sizes per age and temperature are stated in Table 1. Semi-transparent points show raw data. Warm-and cold-acclimation data were collected in two separate studies, as detailed in the main text.

## Discussion

By raising Japanese quail in heatwave-like conditions, we found that birds growing up in warm post-hatch environments were more equipped to counter high air temperature as juveniles compared to control birds. In line with our predictions, warm-reared birds had lower metabolic rate than control birds, and so could dissipate more of their metabolic heat production (MHP) by evaporation compared to control birds when exposed to hot air (i.e., the ratio between evaporative heat loss (EHL) and MHP was higher). Thus, birds growing up warm are more likely to handle heat better in the short term, because evaporative cooling capacity is crucial for heat tolerance (McKechnie and Wolf, 2019; McKechnie et al., 2016). A similar result was obtained in a heat acclimation study on adult zebra finches (*Taeniopygia guttata*) (Wojciechowski et al., 2020), but our data contrast recent work where a postnatal heat wave in zebra finches did not affect MHP or EWL close to fledging (Ton et al., 2021). However, the zebra finches studied by Ton and colleagues had lower *T*_b_ than control birds both at fledging and as juveniles more than a month after the birds had fledged and the experimental heatwave had ended (Ton et al., 2021). In contrast to this, we found no effects on *T*_b_ at any age, and there was also no difference in MHP or evaporative cooling capacity at 8 weeks, when the Warm study birds had been kept in common garden for more than one month. This indicates that evaporative cooling capacity develops faster when birds grow up in warmer conditions, but that the ambient temperature stimulus in this study did not cause permanent effects on thermal and hygric physiology.

We found no evidence that cold-acclimated birds improved their ability of countering low air temperatures as juveniles because growing up cold did not affect MHP, evaporative cooling capacity or *T*_b_. This contrasts the often-observed association between low environmental temperature and increased metabolic rate (reviewed by Swanson and Vézina, 2015). For example, Petit et al, (2013) found a tendency for higher basal metabolic rate (BMR) during the coldest winter months. In line with this, birds measured during a colder winter had higher BMR than those sampled during a milder winter (Dutenhoffer and Swanson, 1996). Since environmental temperature in these studies was well below that in ours, it can be speculated that ambient temperature in the Cold study was not low enough to cause changes in MHP. The aerobic capacity model of endothermy posits that there is a positive correlation between basal (BMR) and maximal heat production (summit metabolic rate; *M*_sum_) (Bennett and Ruben, 1979). However, studies found that this is not always the case (Petit et al., 2013; Swanson et al., 2012), possibly because BMR and *M*_sum_ are driven by different organs (Barceló et al., 2017; Petit et al., 2013). Thus, lack of effects on MHP in our cold birds does not preclude the possibility that cold acclimation contributed changes in *M*_sum_. This should be addressed in future studies.

In contrast to our prediction, cold acclimation did not come at a cost to EWL upon exposure to high air temperature and, in the absence of the expected increase in MHP, also did not truncate evaporative cooling capacity. Thus, we found no evidence for a trade-off between heat-and cold-acclimation at the level of thermoregulation. While such a trade-off might be perceived as intuitive and sometimes occurs at the organismal level (Shou et al., 2022), it need not be expected on regulatory grounds where thermogenesis and thermolysis are not physiologically linked. Nonetheless, we found that cold-acclimated birds had higher baseline (i.e., non-heat-induced) EWL than control birds (Fig. 3D), which is similar to studies of cold-acclimation in passerines (Wang et al., 2019; Williams and Tieleman 2000) at all but the highest test temperature. However, this response was accompanied by elevated MHP in the studies by Wang et al. (2019) and Williams and Tieleman (2000), suggesting that higher EWL could have been (at least in part) explained by increased respiratory EWL. The mechanism explaining higher baseline evaporation in cold-acclimated birds in our study is unclear. Specifically, we found neither strong support for a cold-induced increase in MHP that could be suggestive of increased respiratory water loss, nor a remaining effect on EWL in the heat (i.e., at 40°C), which should be expected if there were underlying differences in cutaneous evaporation between the cold and control birds.

Even though ecogeographical rules suggest body size could be expected to decrease and increase in the warmth and cold, respectively, we found no effect of developmental ambient temperature on body mass and wing length in our study. It is possible that environmental temperature-dependent constraints on development are not much of an issue when there is unrestricted access to food and water. This is different from the effects of high or low environmental temperature in the wild, where heatwave-conditions are often associated with reduced food availability and drought (Kingsolver and Woods, 1997; Smith, 1982; reviewed by Cunningham et al., 2021) and cold snap-conditions can result in lower food abundance (Shipley et al., 2020; Garett et al., 2022). Moreover, while many studies have shown that high environmental temperatures cause decreases in body size (e.g., Salewski et al., 2010; Andrew et al., 2017; Andreasson et al., 2018), there are several exceptions to this pattern (Salewski et al., 2010; Siepielski et al., 2019; Ton et al., 2021). Thus, emergence of small or large body size to match environmental temperatures is not a general response and may be driven more strongly by efficiency of energy use than thermal advantage per se (reviewed by Tabh and Nord, 2023). Nonetheless, Burness et al. (2013) and Andrew et al. (2017) report on reduced size in heat-challenged captive birds with *ad libitum* access to food and water. Moreover, broilers raised for meat production gain less body mass and eat less food, and have higher mortality, when exposed to both acute (Zaboli et al., 2017; Arjona et al., 1988) and chronic (Imik et al., 2012; Sohail et al., 2012; Ranjan et al., 2019) heat stress. Perhaps the difference between the control and warm or cold treatment in our study was too small, the warm treatment temperature too low, and the cold treatment temperature too high, to cause any irreversible changes on body size. Alternatively, it is possible that other selection pressures for large or small body size, such as competitive ability in males or egg size in females, were stronger than selection for any size-related increase in dry heat flux in the warmth or a decrease thereof in the cold; demands for which would be more easily accommodated by physiological flexibility within a generation. Future studies should address if developmental environmental temperature affect size or length of the appendages. According to Allen’s rule (Allen, 1877), appendages such as the legs and bills of birds or the tails and ears of mammals, should be longer in warmer environmental temperatures (cf. Demicka and Caputa, 1993) and shorter in the cold environmental temperatures. Since bird bills and legs are both richly vascularised and uninsulated, changes in local circulation can quickly either increase or decrease heat flux (Scholander et al., 1950; Tattersall et al., 2009). Thus, appendage size is likely aiding thermoregulation by the whole bird much more than any change in size per se, though broad generalisations of the biological meaning of such effects are probably naïve (Tabh and Nord, 2023).

## Conclusions

The environmental matching hypothesis predicts that animals growing up in warmer environments should be better at handling warm environmental temperatures as adults, which is suggested in some studies (e.g., Demicka and Caputa, 1993; Stawski and Geiser, 2020; reviewed by Nord and Giroud, 2020), and vice versa for animals growing up in the cold. In other studies, silver spoon effects appear to be more common, reflected by reduced thermoregulatory performance or constrained growth after development in thermally challenging conditions (reviewed by Andreasson et al., 2020; Nord and Giroud, 2020). We found support for neither hypothesis. However, it is important to note that we did not measure maximum heat and cold tolerance, which might be more telling of thermal acclimation than submaximal physiological rates. Such measurements could also be more informative when extrapolating laboratory results to free-ranging animals because, in the wild, environmental temperatures vary by several tens of degrees between different microhabitats, for example in relation to vegetation and topography (Carroll et al., 2016; Cunningham et al., 2015; Cunningham et al., 2021). Knowledge of temperature tolerances will inform on which animals are better able to exploit resources spread across such thermal landscapes. Future studies should, therefore, address if a longer heatwave-or cold snap period, or a more severe environmental temperature treatment, cause long-term or irreversible effects. It can be speculated that effects of developmental temperature on thermoregulation could be caused by changes in levels of hormones that regulate energy metabolism. Thyroid hormones, regulated via the hypothalamus-pituitary-thyroid (HPT) axis, are positively linked to thermogenesis (reviewed by Ruuskanen et al., 2021). Studies have found that the HPT axis is sensitive to developmental heat stimulation, such that broilers exposed to a heat stress in ovo have lower plasma concentration of triiodothyronine (T_3_) in adolescence, especially during a heat challenge (Piestun et al., 2008). Moreover, the glucocorticoid hormone corticosterone, regulated via the hypothalamus-pituitary-adrenal (HPA) axis, is the major stress hormone in birds and necessary for keeping homeostasis (reviewed by Ruuskanen et al., 2021). It increased in response to both heat-(Piestun et al., 2008) and cold (Bize et al., 2010) exposure. Finally, when thermoregulation depend on evaporative cooling such as in the highest measurement temperature in our study, arginine vasotocin (AVT) could be an important candidate hormone to study due to its role in water balance (Goldstein, 2006). Studies have found that AVT increases in response to dehydration (Robinzon et al., 1990) and that administration of exogenous AVT leads to fluid retention (Takahashi et al., 1995) and reduces *T*_b_ (John and George, 1992; Hassinen et al., 1999). We suggest that future work investigate the mechanisms mediating developmental thermal acclimation, even when these effects are reversible like in this study.

## Ethics

Ethical approval for the experiment was granted by the Malmö/Lund Animal Ethics Committee (permit no. 9246-19).

### LIST OF ABBREVIATIONS

AVT: arginine vasotocin
BMR: basal metabolic rate
EHL: evaporative heat loss
EHL/MHP: evaporative cooling capacity, defined as the ratio between evaporative heat loss and metabolic heat production
EWL: evaporative water loss
HPA: hypothalamus-pituitary-adrenal
HPT: hypothalamus-pituitary-thyroid
MHP: metabolic heat production
*M*_sum_: summit metabolic rate
PIT: passive integrated transponder
STPD: standard temperature and pressure, dry
T_3_: triiodothyronine
*T*_b_: body temperature

## ACKNOWLEDGEMENTS

Lars Råberg, Fredrik Andreasson and three anonymous reviewers provided helpful comments that improved previous drafts of the manuscript. We thank Camilla Björklöv and Agnieszka Czopek for assistance in caring for the birds, Lars Fredriksson for all the help with building pens and chambers and Cecilia Thomasson for providing equipment.

